# Increased and decreased superficial white matter structural connectivity in schizophrenia and bipolar disorder

**DOI:** 10.1101/473686

**Authors:** Ellen Ji, Pamela Guevara, Miguel Guevara, Antoine Grigis, Nicole Labra, Samuel Sarrazin, Nora Hamdani, Frank Bellivier, Marine Delavest, Marion Leboyer, Ryad Tamouza, Cyril Poupon, Jean-François Mangin, Josselin Houenou

## Abstract

Schizophrenia (SZ) and bipolar disorder (BD) are often conceptualized as “disconnection syndromes”, with substantial evidence of abnormalities in deep white matter tracts, forming the substrates of long-range connectivity, seen in both disorders. However, the study of superficial white matter (SWM) U-shaped short-range tracts remained challenging until recently, although findings from post-mortem studies suggest they are likely integral components of SZ and BD neuropathology. This diffusion weighted imaging (DWI) study aimed to investigate SWM microstructure *in vivo* in both SZ and BD for the first time. We performed whole brain tractography in 31 people with SZ, 32 people with BD and 54 controls using BrainVISA and Connectomist 2.0. Segmentation and labelling of SWM tracts were performed using a novel, comprehensive U-fiber atlas. Analysis of covariances yielded significant generalized fractional anisotropy (gFA) differences for 17 SWM bundles in frontal, parietal and temporal cortices. Post hoc analyses showed gFA reductions in both patient groups as compared with controls in bundles connecting regions involved in language processing, mood regulation, working memory and motor function (pars opercularis, insula, anterior cingulate, precentral gyrus). We also found increased gFA in SZ patients in areas overlapping the default mode network (inferior parietal, middle temporal, precuneus), supporting functional hyperconnectivity of this network evidenced in SZ. We thus illustrate that short U-fibers are vulnerable to the pathological processes in major psychiatric illnesses, encouraging improved understanding of their anatomy and function.

## Introduction

Theories of disconnection among neurological disorders emerged over a century ago ^1^ and have been followed by the conceptualization of schizophrenia (SZ) as a “disconnection syndrome” ^2^. Indeed, convergent lines of evidence indicate that the microstructure of white matter (WM) tracts is compromised in at least some people diagnosed with SZ and these findings have been extended to bipolar disorder (BD) ^3, 4^. Whereas deep WM tracts form the substrates of long-range connectivity, superficial white matter (SWM) is comprised of short association bundles (often referred to as “U-fibers” due to their appearance) and lies just beneath the grey matter tissue of the cortex, mediating local connectivity between adjacent cortical gyri. This shallow layer of WM is the last area to myelinate, enabling high plasticity capabilities but also consequently tremendous vulnerability ^5^. An increase of interstitial white matter neuron (IWMN) density has been found in the SWM of SZ patients ^6–9^ in neuropathological studies, which may be correlated with an interneuron deficit ^6^, a strongly supported theory of SZ pathology. Yet, while studies have examined SWM in postmortem brain tissue of patients, very few studies using *in vivo* methods have been performed.

Diffusion-weighted imaging (DWI) is a non-invasive technique that tracks the diffusion properties of water through brain tissue, providing a sensitive measure of white matter integrity. The most commonly assessed scalar value in DWI studies is fractional anisotropy (FA), which reflects the extent to which water diffusion is directionally restricted along a single axis, known as anisotropic diffusion. Reliable measurements of FA rely on robust estimation of the diffusion tensor, which is associated with the number of diffusion-encoding gradient directions (NDGD) ^10, 11^: an increased NDGD yields a more accurate diffusion tensor calculation. Human and animal studies have shown that FA is correlated with myelination, fiber coherence and number of axons ^12–16^. In SZ, nearly all DWI studies have focused on long-range deep WM tracts ^17–20^ likely because SWM is more complex and smaller in size, has high inter-subject variability and a tailored SWM atlas was only developed last year ^21^. To our knowledge, only two studies have examined SWM *in vivo* in SZ ^22, 23^. Without an atlas of discrete SWM tracts and as their NDGDs were 6 and 23, the specificity of these findings may be difficult to replicate with adequate detail and accuracy. Moreover, tractography methods were not employed despite the advantages of tractography over some voxel-based studies. Indeed, their differential findings of SWM FA reductions in people with SZ primarily in the left frontal lobe ^22^ versus SWM FA reductions in the temporal and occipital regions ^23^ fit with the disconnectivity hypothesis but warrant further investigation. Two studies have studied SWM in BD ^24, 25^. Cabeen et al. used DWI voxel-wised methods but did not find a significant main group effect between BD and control groups ^24^ and a very recent study by Zhang et al. used a combination of tractography (to define a SWM mask) and TBSS (for statistical analyses) and found decreased FA proximal to regions related to the emotion dysregulation in BD ^25^.

There is considerable overlap in the neurobiological features, clinical symptoms and genetic vulnerability of SZ and BD ^26^. While still currently classified as two separate mental disorders, their similarities and unsatisfactory treatment response rate underscore the urgent importance of better understanding their biological profiles and advancing knowledge. The present DWI study examined SWM *in vivo* using tractography with a specific atlas, and a high NDGD of 60, in people diagnosed with SZ, BD and controls. Our primary aim was to explore anatomically-delineated SWM tracts using DWI-based tractography and a U-fibers atlas ^23, 27^ to determine abnormalities that are common and disease-specific to SZ and BD.

## Methods

### Participants

Patients with a diagnosis of schizophrenia (23 men and 8 women) or bipolar disorder (21 men and 11 women) were recruited from the psychiatry departments of Mondor University Hospital, Créteil, France and Fernand-Widal Lariboisière University Hospital, Paris, France. Healthy adults (23 men and 31 women) were recruited from the general population through advertising as a comparison group. All participants were between the ages of 15 and 55 years. Diagnostic and exclusion criteria, and clinical and cognitive measures are described in the Supplement Materials. The study protocol was approved by the local ethics committee (CPP Ile de France IX). All subjects provided written informed consent after receiving a complete description of the study and prior to participation.

### Image acquisition

Structural MRI scans were acquired using a 3-Tesla Siemens Magnetom Tim Trio scanner with 12 channel head coil at NeuroSpin, CEA, Saclay, France. Each participant received a T1-weighted high-resolution anatomical scan with the following parameters: repetition time (TR) = 2300 ms; echo time (TE) = 2.98 ms; FOV = 256 mm^2^; voxel size = 1 × 1 × 1.1 mm^3^; 180 slices and a shared diffusion weighted sequence along 60 directions with voxel size = 2 × 2 × 2 mm^3^; b = 1400 s/mm plus one image in which b = 0.

### Data Processing

We performed whole brain deterministic tractography and data processing using a validated processing pipeline ^28, 29^. T1-weighted and DW images were processed using BrainVISA 4.2 (http://www.brainvisa.info) and Connectomist 2.0, respectively. Data were assessed for movement, susceptibility and noise artefacts for both T1 and DW acquisitions. For DW data, an orientation distribution function at each voxel was computed, containing information about the angular profile of diffusion within each voxel, which in turn allows detection of principal directions of diffusivity similar to the main eigenvector of DTI models. We used the Q-ball imaging (QBI) model which better models diffusivity in WM areas of complex architecture and organisation (i.e. crossing fibers) ^30^ than the classical diffusion tensor model. The mean generalized fractional anisotropy (gFA) from all the computed orientation distribution functions was evaluated ^30^. Whole brain tractography and tractogram segmentation are detailed in the Supplementary Methods.

### Statistical analyses

All data analyses were performed with the α value set to 0.05. Antipsychotic dose was converted to mean daily olanzapine (OLZ) equivalent dose based on standard guidelines ^31^. Demographic variables, mean current daily OLZ equivalent dose and premorbid IQ were compared among groups using univariate analyses of variance or *X*^2^-tests, as appropriate.

To determine whether there were gFA differences among the patient groups and healthy controls, multivariate analyses of covariances were applied to gFA of bundles with diagnosis (3 levels: control, SZ, BD) as the between-group factor and age and sex as covariates. False-positive results related to multiple comparisons were controlled using the Benjamini-Hochberg false discovery rate (FDR) method ^32^. Effect size calculations were measured as partial eta squared (η^2^). Follow up pairwise comparisons were then conducted with FDR corrections. For bundles identified as having significantly different gFA values between BD patients and controls, we compared mean gFA between subgroups of BD patients with and without a history of psychotic features. To examine whether SWM microstructure may change over the course of the illness, we performed correlations between gFA and illness duration, separately in patients with SZ and BD, across all bundles while controlling for age, as illness duration and age are collinear.

We assessed the potential influence of antipsychotic medication on SWM integrity by performing Pearson’s correlations between OLZ equivalent scores and gFA for patients (SZ and BD) taking antipsychotics. To investigate whether gFA was associated with symptom severity in patients, we performed Pearson’s correlations between symptom severity scores - as measured by PANSS positive, negative, general and total - and gFA of bundles that were significantly different following post hoc analyses, separately in patients with SZ and BD.

## Results

Demographic and clinical characteristics of participants are presented in Table 1. Participants with SZ showed mild to moderate symptom severity based on the PANSS scores. There was no significant difference in age between the groups. There was a significant difference in sex ratios in which there was a greater proportion of female participants in the control group relative to patient groups. There was an expected difference in education in which healthy controls received more years of education relative to both patient groups.

**Table 1.**
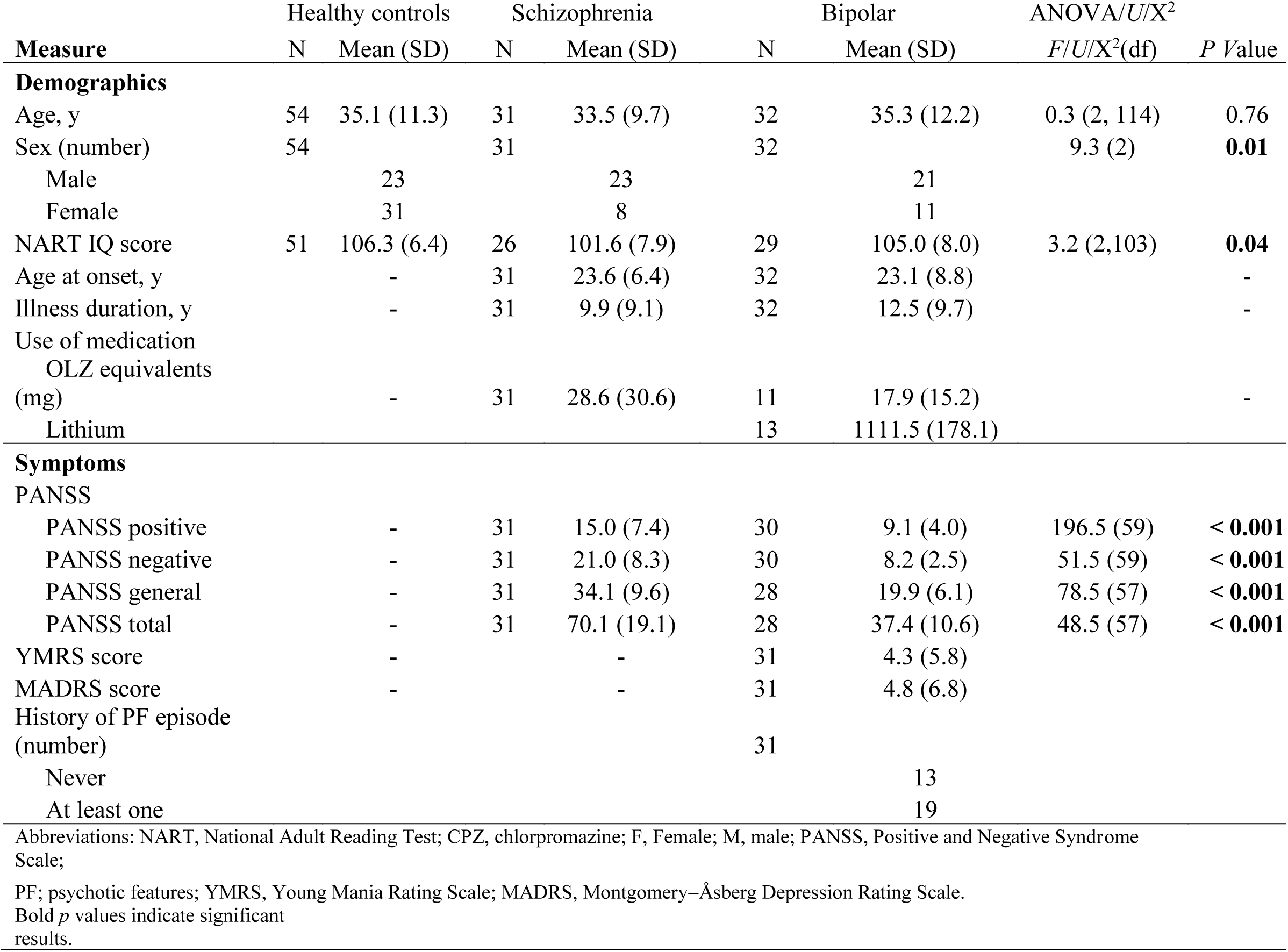
Demographic variables and clinical characteristics of the whole sample.

| Measure | Healthy controls | | Schizophrenia | | Bipolar | | ANOVA/$U/X^2$ | |
| --- | --- | --- | --- | --- | --- | --- | --- | --- |
| | N | Mean (SD) | N | Mean (SD) | N | Mean (SD) | $F/U/X^2(df)$ | $P$ Value |
| **Demographics** |  |  |  |  |  |  |  |  |
| Age, y | 54 | 35.1 (11.3) | 31 | 33.5 (9.7) | 32 | 35.3 (12.2) | 0.3 (2, 114) | 0.76 |
| Sex (number) | 54 |  | 31 |  | 32 |  | 9.3 (2) | **0.01** |
| Male |  | 23 |  | 23 |  | 21 |  |  |
| Female |  | 31 |  | 8 |  | 11 |  |  |
| NART IQ score | 51 | 106.3 (6.4) | 26 | 101.6 (7.9) | 29 | 105.0 (8.0) | 3.2 (2,103) | **0.04** |
| Age at onset, y |  | - | 31 | 23.6 (6.4) | 32 | 23.1 (8.8) |  | - |
| Illness duration, y |  | - | 31 | 9.9 (9.1) | 32 | 12.5 (9.7) |  | - |
| Use of medication |  |  |  |  |  |  |  |  |
| OLZ equivalents |  |  |  |  |  |  |  |  |
| (mg) |  | - | 31 | 28.6 (30.6) | 11 | 17.9 (15.2) |  | - |
| Lithium |  |  |  |  | 13 | 1111.5 (178.1) |  |  |
| **Symptoms** |  |  |  |  |  |  |  |  |
| PANSS |  |  |  |  |  |  |  |  |
| PANSS positive |  | - | 31 | 15.0 (7.4) | 30 | 9.1 (4.0) | 196.5 (59) | < **0.001** |
| PANSS negative |  | - | 31 | 21.0 (8.3) | 30 | 8.2 (2.5) | 51.5 (59) | < **0.001** |
| PANSS general |  | - | 31 | 34.1 (9.6) | 28 | 19.9 (6.1) | 78.5 (57) | < **0.001** |
| PANSS total |  | - | 31 | 70.1 (19.1) | 28 | 37.4 (10.6) | 48.5 (57) | < **0.001** |
| YMRS score |  | - |  | - | 31 | 4.3 (5.8) |  |  |
| MADRS score |  | - |  | - | 31 | 4.8 (6.8) |  |  |
| History of PF episode |  |  |  |  |  |  |  |  |
| (number) |  |  |  |  | 31 |  |  |  |
| Never |  |  |  |  |  | 13 |  |  |
| At least one |  |  |  |  |  | 19 |  |  |
| Abbreviations: NART, National Adult Reading Test; CPZ, chlorpromazine; F, Female; M, male; PANSS, Positive and Negative Syndrome Scale; |  |  |  |  |  |  |  |  |
| PF; psychotic features; YMRS, Young Mania Rating Scale; MADRS, Montgomery–Åsberg Depression Rating Scale. |  |  |  |  |  |  |  |  |
| Bold $p$ values indicate significant results. | | | | | | | | |

### Imaging

Out of the 100 bundles from the atlas used for segmentation and labelling, 65 were successfully reconstructed and thus stable in all three study groups. Multivariate analyses of covariances determined that gFA of 17 out of the 65 stable bundles were significantly different among groups (Table 3). Post hoc analyses (Supplementary Table 2, Figure 1) revealed significant reductions in mean gFA in SZ patients as compared with controls in 13 of the 17 bundles including the left CAC-PrC, bilateral CMF-PrC, left MOF-ST, left Op-Ins, bilateral Op-PrC, left PoC-Ins, right IP-MT, right LOF-RMF, right PoC-PrC, right PoC-SM and right RMF-SF. There were also 13 bundles in which BD patients showed lower gFA compared with controls which included the left CAC-PrCu, bilateral CMF-PrC, bilateral IP-MT, left Op-Ins, left PoC-PrC, left PoC-Ins, left PoCi-PrCu, right MOF-ST, right PoC-PrC, right PoC-SM and right RMF-SF. The direct comparison between the two patient groups revealed that patients with SZ had significantly lower gFA in the left CAC-PrCu, left MOF-ST, right CMF-PrC, right LOF-RMF, right Op-PrC, right PoC-PrC and right RMF-SF, and significantly greater gFA in the bilateral IP-MT, left Op-PrC, left PoCi-PrC, left PoC-SM compared to BD patients. Controls had decreased gFA in the left IP-MT, left PoCi-PrCu, left PoC-SM compared with SZ patients and in three bundles including the left PoC-SM, right LOF-RMF and right Op-PrC compared with BD patients. According to Cohen’s criteria (partial eta squared values of 0.01, 0.06, and 0.14 correspond to small, medium, and large effect sizes, respectively) ^33^, 12 out of 17 bundles that were significant among groups had large effect sizes while the remaining 5 bundles were within 0.02 of a large effect size.

**Figure 1.**
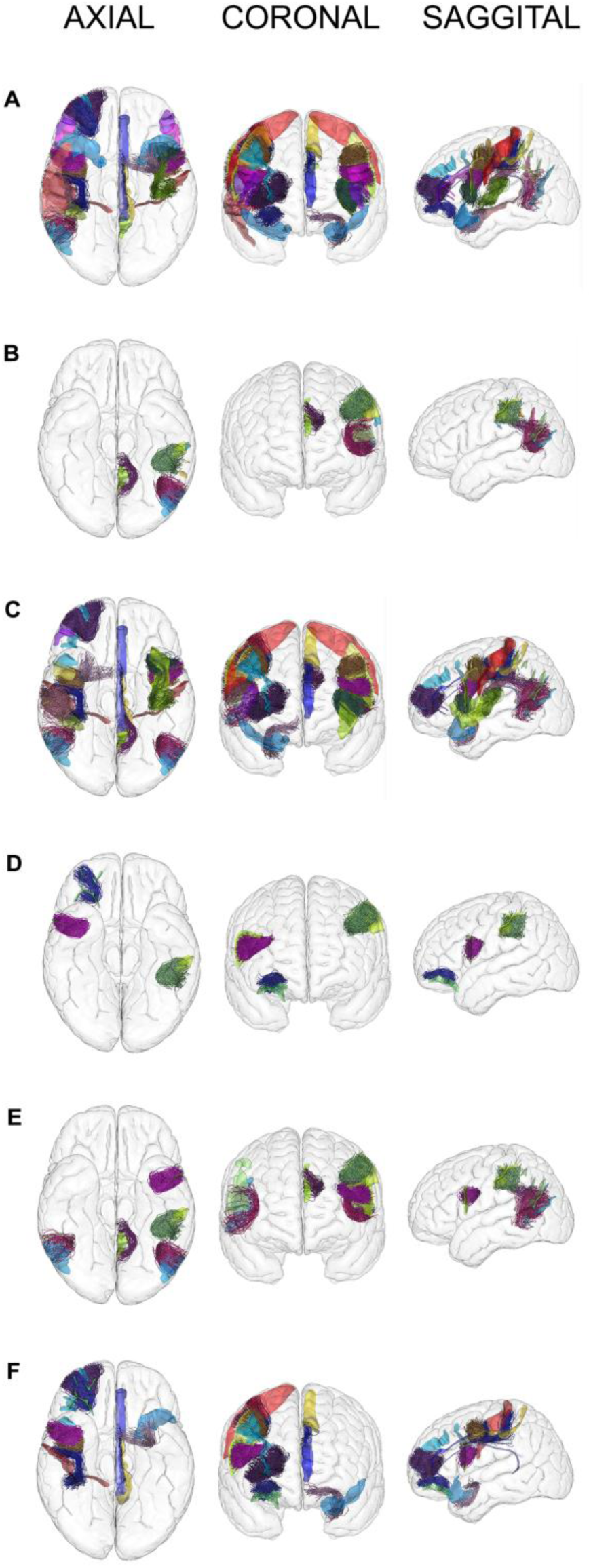
Axial, coronal and sagittal views of post hoc results (mean gFA) demonstrating differences between groups. Images are displayed using radiological convention. (A) gFA Controls > Schizophrenia (B) gFA Schizophrenia > Controls (C) gFA Controls > Bipolar (D) gFA Bipolar > Controls (E) gFA Schizophrenia > Bipolar (F) gFA Bipolar > Schizophrenia

**Table 2.**
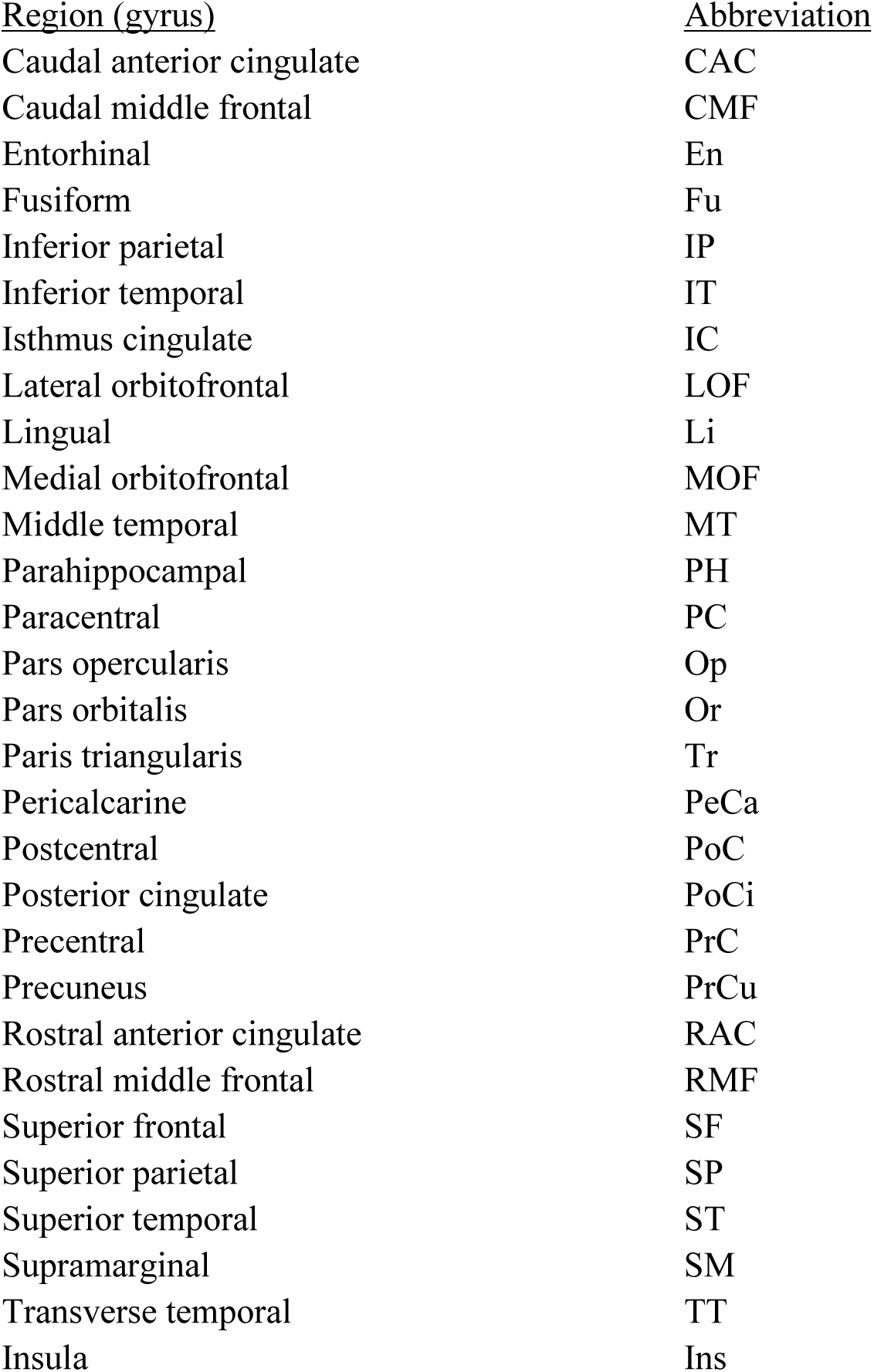
Anatomical abbreviations

| <u>Region (gyrus)</u> | <u>Abbreviation</u> |
| --- | --- |
| Caudal anterior cingulate | CAC |
| Caudal middle frontal | CMF |
| Entorhinal | En |
| Fusiform | Fu |
| Inferior parietal | IP |
| Inferior temporal | IT |
| Isthmus cingulate | IC |
| Lateral orbitofrontal | LOF |
| Lingual | Li |
| Medial orbitofrontal | MOF |
| Middle temporal | MT |
| Parahippocampal | PH |
| Paracentral | PC |
| Pars opercularis | Op |
| Pars orbitalis | Or |
| Pars triangularis | Tr |
| Pericalcarine | PeCa |
| Postcentral | PoC |
| Posterior cingulate | PoCi |
| Precentral | PrC |
| Precuneus | PrCu |
| Rostral anterior cingulate | RAC |
| Rostral middle frontal | RMF |
| Superior frontal | SF |
| Superior parietal | SP |
| Superior temporal | ST |
| Supramarginal | SM |
| Transverse temporal | TT |
| Insula | Ins |

**Table 3.** Between-group gFA Comparisons of SWM Bundles

| Bundle | Hemisphere | F (df) | p Value | p Value<br>for FDR<br>Adjusted | Partial<br>$\eta^2$ |
| --- | --- | --- | --- | --- | --- |
| CAC - PrCu | L | 7.7 (4, 112) | <0.001 | **<0.001<sup>a,b,c</sup>** | 0.22 |
| CMF - PrC | L | 1.0 (4, 112) | 0.4 | 0.46 | 0.03 |
| CMF - PrC_2 | L | 3.9 (4, 112) | 0.005 | **0.02<sup>a,b</sup>** | 0.12 |
| CMF - RMF | L | 0.6 (4, 112) | 0.63 | 0.65 | 0.02 |
| CMF - SF | L | 1.3 (4, 112) | 0.28 | 0.35 | 0.04 |
| IC - PrCu | L | 0.7 (4, 112) | 0.56 | 0.60 | 0.03 |
| IP - LOF | L | 2.3 (4, 112) | 0.06 | 0.14 | 0.08 |
| IP - MT | L | 5.7 (4, 112) | <0.001 | **<0.001<sup>a,b,c</sup>** | 0.17 |
| IP - SP | L | 0.3 (4, 112) | 0.85 | 0.85 | 0.01 |
| IT - MT | L | 1.3 (4, 112) | 0.26 | 0.34 | 0.05 |
| LOF - Or | L | 1.7 (4, 112) | 0.15 | 0.23 | 0.06 |
| LOF - RMF | L | 2.8 (4, 112) | 0.03 | 0.09 | 0.09 |
| LOF - RMF_2 | L | 2.0 (4, 112) | 0.10 | 0.19 | 0.07 |
| LOF - ST | L | 2.7 (4, 112) | 0.03 | 0.09 | 0.09 |
| MOF - ST | L | 4.0 (4, 112) | 0.004 | **0.02<sup>a,c</sup>** | 0.13 |
| Op - Ins | L | 4.8 (4, 112) | 0.001 | **0.006<sup>a,b</sup>** | 0.15 |
| Op - PrC | L | 5.4 (4, 112) | 0.001 | **0.006<sup>a,b,c</sup>** | 0.17 |
| Op - SF | L | 0.4 (4, 112) | 0.8 | 0.81 | 0.01 |
| Or - Ins | L | 1.6 (4, 112) | 0.17 | 0.25 | 0.06 |
| PoC - Ins | L | 4.0 (4, 112) | 0.004 | **0.02<sup>a,b</sup>** | 0.13 |
| PoCi - PrCu | L | 5.2 (4, 112) | 0.001 | **0.006<sup>a,b,c</sup>** | 0.16 |
| PoCi - RAC | L | 2.4 (4, 112) | 0.06 | 0.14 | 0.08 |
| PoCi - SF | L | 1.14 (4, 112) | 0.34 | 0.40 | 0.04 |
| PoC - PrC | L | 1.8 (4, 112) | 0.13 | 0.22 | 0.06 |
| PoC - PrC_2 | L | 2.1 (4, 112) | 0.09 | 0.17 | 0.07 |
| PoC - PrC_3 | L | 1.5 (4, 112) | 0.19 | 0.27 | 0.05 |
| PoC - PrC_4 | L | 1.6 (4, 112) | 0.17 | 0.25 | 0.06 |
| PoC - SM | L | 1.1 (4, 112) | 0.37 | 0.42 | 0.04 |
| PoC - SM_2 | L | 5.3 (4, 112) | 0.001 | **0.006<sup>a,b,c</sup>** | 0.16 |
| PrC - Ins | L | 2.3 (4, 112) | 0.06 | 0.14 | 0.08 |
| RMF - SF | L | 2.3 (4, 112) | 0.06 | 0.14 | 0.08 |
| SP - SM | L | 1.8 (4, 112) | 0.14 | 0.23 | 0.06 |
| ST - TR | L | 1.7 (4, 112) | 0.16 | 0.24 | 0.06 |
| Pr - Ins | L | 1.4 (4, 112) | 0.23 | 0.32 | 0.05 |
| CAC - PrCu | R | 2.3 (4, 112) | 0.07 | 0.14 | 0.08 |
| CMF - PrC | R | 1.9 (4, 112) | 0.11 | 0.19 | 0.07 |
| CMF - PrC_2 | R | 4.6 (4, 112) | 0.002 | **0.01<sup>a,b,c</sup>** | 0.14 |
| CMF - SF | R | 1.4 (4, 112) | 0.25 | 0.33 | 0.05 |
| CMF - SF_2 | R | 2.5 (4, 112) | 0.05 | 0.13 | 0.08 |
| IC - PrCu | R | 1.7 (4, 112) | 0.15 | 0.23 | 0.06 |
| IP - IT | R | 2.5 (4, 112) | 0.05 | 0.13 | 0.08 |
| IP - MT | R | 8.8 (4, 112) | <0.001 | **<0.001<sup>a,b,c</sup>** | 0.24 |
| IP - SP | R | 1.2 (4, 112) | 0.31 | 0.37 | 0.04 |
| IT - MT | R | 1.1 (4, 112) | 0.35 | 0.40 | 0.04 |
| LOF - MOF | R | 3.0 (4, 112) | 0.02 | 0.08 | 0.10 |
| LOF - RMF | R | 4.2 (4, 112) | 0.003 | **0.02<sup>a,b,c</sup>** | 0.13 |
| LOF - ST | R | 1.6 (4, 112) | 0.19 | 0.27 | 0.05 |
| MOF - ST | R | 3.9 (4, 112) | 0.005 | **0.02<sup>b</sup>** | 0.12 |
| MT - ST | R | 2.2 (4, 112) | 0.08 | 0.16 | 0.07 |
| Op - Ins | R | 2.8 (4, 112) | 0.03 | 0.09 | 0.09 |
| Op - PrC | R | 6.8 (4, 112) | <0.001 | **<0.001<sup>a,b,c</sup>** | 0.20 |
| Or - Ins | R | 2.8 (4, 112) | 0.03 | 0.09 | 0.09 |
| PoCi - PrCu | R | 0.9 (4, 112) | 0.46 | 0.49 | 0.03 |
| PoCi - RAC | R | 2.3 (4, 112) | 0.07 | 0.14 | 0.08 |
| PoC - PrC | R | 1.0 (4, 112) | 0.41 | 0.46 | 0.03 |
| PoC - PrC_2 | R | 2.0 (4, 112) | 0.10 | 0.19 | 0.07 |
| PoC - PrC_3 | R | 5.1 (4, 112) | 0.001 | **0.006<sup>a,b,c</sup>** | 0.16 |
| PoC - SM | R | 5.0 (4, 112) | 0.001 | **0.006<sup>a,b</sup>** | 0.15 |
| PoC - SP | R | 1.9 (4, 112) | 0.11 | 0.19 | 0.06 |
| PrC - Ins | R | 1.4 (4, 112) | 0.25 | 0.33 | 0.05 |
| RAC - SF | R | 0.7 (4, 112) | 0.58 | 0.60 | 0.03 |
| RMF - SF | R | 6.3 (4, 112) | <0.001 | **<0.001<sup>a,b,c</sup>** | 0.18 |
| SP - SM | R | 1.2 (4, 112) | 0.30 | 0.37 | 0.04 |
| ST - TT | R | 2.1 (4, 112) | 0.09 | 0.17 | 0.07 |
| Tr - Ins | R | 2.2 (4, 112) | 0.08 | 0.16 | 0.07 |

Abbreviation: FDR, false discovery rate.
Bold $p$ values indicate significant ANCOVA results.
<sup>a</sup> Healthy controls significantly different from schizophrenia group ( $p \leq 0.05$ , FDR adjusted).
<sup>b</sup> Healthy controls significantly different from bipolar group ( $p \leq 0.05$ , FDR adjusted).
<sup>c</sup> Schizophrenia group significantly different from bipolar group ( $p \leq 0.05$ , FDR adjusted).

### Clinical correlations

Nineteen out of 32 BD patients (59%) had a history of psychotic features. We did not observe any differences in mean gFA in any ROIs between BD patients with and without a history of psychotic features (Supplementary Table 3). Mean gFA was not associated with illness duration in any bundle (Supplementary Table 4). Negative correlations were observed between OLZ equivalent scores and gFA in nearly one-fifth of bundles, with six bundles surviving correction for multiple comparisons in patients taking antipsychotic mediation (Supplementary Table 5).

There was a positive correlation between gFA in one bundle (left PoC-SM) and negative symptoms in people with SZ. There was no association between symptom severity and gFA in any other bundle in people with SZ or BD (Supplementary Table 6).

## Discussion

In the first study to examine SWM in both SZ and BD *in vivo* including a direct comparison between the two disorders, we found significant differences in gFA throughout frontal, parietal and temporal cortices of the brain, after correction for multiple comparisons. Only two studies have previously investigated SWM in SZ, and two other studies in BD, using diffusion imaging. Replication of results is essential to ensure that biases and confounders are not driving the results ^34^. We were able to replicate findings by Phillips and colleagues ^23^ of decreased FA in SZ in the left temporal lobe in people with SZ and findings by Nazeri et al. ^27^ of lower FA in the left PrCu, left inferior frontal gyrus (Op) and left PrC in people with SZ. In BD patients, we, like others ^25^, found reduced FA in the left CMF-PrC. In extending these findings, our study detected differences in several additional regions including localized increased gFA in patients, which we attribute to the specific methods we used (advanced QBI model, tractography method and tailored SWM atlas).

The overall pattern common to both SZ and BD was widespread decreased gFA. Shared abnormalities in gFA in SZ and BD relative to controls reflect impairments common to both disorders. Moreover, the shared abnormalities support common disease-related genetic liability and is in line with findings that SWM FA varies in accordance with relatedness in unaffected relatives of patients with SZ ^23^. Patients showed reduced gFA in the bilateral CMF-PrC and right PoC-PrC forming connections to/from the primary motor cortex, which may represent shared impaired motor function ^35, 36^. Likewise, our finding of decreased gFA in patients in the left Op-PrC and Op-Ins, which contribute to speech production and processing including Broca’s area, denote linguistic impairments ^37–41^. Our finding in areas involved in executive functions and working memory ^42^, CAC-PrCu and RMF-SF, may be related to the widespread cognitive impairments in SZ that are milder in BD ^43^ as the decrease in gFA was even more pronounced in people with SZ relative to people with BD. In support of this, previous reports have found SWM FA of the left frontal lobe to predict attention and working memory performance in healthy people, but not in patients who showed decreases in FA [23]. SZ patients also showed significantly decreased gFA compared to controls and BD patients of the left MOF-ST, which has received substantial attention in the investigation of SZ’s neurobiology and is implicated in disinhibited behaviour, cognitive and emotional problems, as well as auditory verbal hallucinations ^44–46^.

The focal differences between SZ and BD patients may evidence lateralized dysfunction where people with SZ tend to exhibit left hemisphere dysfunction to a greater extent than right, whereas the reverse is seen in people with BD ^47^. Indeed, as compared with controls, people with SZ showed increased gFA only in the left hemisphere while people with BD had predominantly increases in the right hemisphere. One explanation for increased gFA in patients is that it may reflect compensatory lateralization. Our observation of decreased gFA in patients in the left Op-PrC, which we believe to be related to impaired speech production and processing, was accompanied by increased gFA in BD patients as compared to both controls and SZ patients in the right Op-PrC. Furthermore, in both patient groups, gFA of the left PoC-SM was increased while its counterpart bundle in the right hemisphere was decreased. The left PoC-SM was also the only bundle correlated with symptom severity and this finding was specific to negative symptoms in the SZ group. However, the directionality of this correlation (positive) was not expected and as there was no other pattern of neural correlates and clinical presentation, it is difficult to comment on its significance. In contrast to an exceedingly simplistic yet prevailing viewpoint that more FA is always superior and in sharp contrast to demyelinating disorders such as multiple sclerosis ^48^, previous research has shown that patients who hear conversing hallucinations have increased FA in interhemispheric auditory fibers compared to patients without this symptom and healthy controls ^49^. Moreover, the severity of hallucinations may be positively correlated with deep WM FA in temporal tracts, such as the arcuate fasciculus and superior longitudinal fasciculus ^50–52^. We observed a pattern of increased gFA in the left IP-MT and PoCi-PrCu in people with SZ, overlapping with the default mode network that has been shown to be functionally hyperconnected in some studies in SZ ^53–55^. This hyperconnectivity is insufficiently suppressed during working memory and is more prominent in cognitively impaired compared to cognitively preserved patients ^56^. Our findings in the PoCi may be linked to psychotic symptoms as structural and functional disturbances here blur the line between internal and external thoughts, thereby assigning self-relevance to unrelated external events ^57^.

Methodological differences between the present study and previous studies may account for dissimilarities in results. Firstly, the four previous neuroimaging studies examining SWM in SZ and BD used either tract-based spatial statistics (TBSS) or surface-based registration, which employ voxel-based methods that cannot resolve the problem of crossing fibers. Moreover, the former method uses an oversimplified skeleton of average FA from the centre of each tract, often resulting in information loss and misregistration, thus only being suited for studying the core of large tracts ^58^. The advanced QBI model applied in this study has the ability to recognise direction of diffusivity in WM areas of intricate architecture ^30^. Secondly, NDGDs of six and 23, in studies by Nazeri et al. ^27^ and Phillips et al. ^23^, respectively, differed considerably from our study’s NDGD of 60. Evidence suggests that the considerably greater number of directions acquired in our study improves the reliability of FA measurements due to increased accuracy/localization of the diffusion tensor, especially in SWM ^10, 11^. Lastly, differences in patients’ age between studies in addition to differences in age between patients and controls within studies is an important methodological consideration due to the maturation of SWM. SZ patients in our study and Nazeri et al.’s study ^27^ had an average age of 34 and 36, respectively. SZ patients in the study by Phillips et al. were below 30, on average, with the oldest patient being only 46. The development of SWM, resembling an inverted U-shaped curve, may be temporally shifted in people with SZ ^59^. While SWM maturity peaks during childhood through adolescence and declines going into adulthood in healthy people, in individuals with SZ SWM maturity may remain lower than healthy people throughout childhood and adolescence and then peaks towards early adulthood, thus there is a period of time during which SWM may be increased in SZ in comparison with healthy people of the same age ^59^. Likewise, Cabeen and colleagues ^24^ studied SWM in children and adolescents between the ages of 8-17 and showed significantly different maturation trajectories between typically developing controls and youths diagnosed with BD in fronto-temporal-striatal connectivity. BD patients in our study and Zhang et al’s study ^25^ had an average of 35 and 26, respectively. Therefore, study differences in age and disease-related maturation trajectories may explain differences in findings among studies.

Categorization based on phenotypes rather than DSM-IV diagnosis may be a more fruitful method to explore the underlying mechanisms of psychiatric disorders. Indeed, mood and psychotic symptoms often coexist in SZ and BD and present different WM profiles ^29^. We and others ^60, 61^ found no differences in FA in any bundle between BD patients with and without a history of psychosis and, moreover, a large-multi site study did not detect differences between SZ and psychotic BD patients ^62^.

However, Sarrazin et al. reported lower gFA in BD patients with a history of psychotic features than those without along the body of the corpus callosum ^29^, suggesting that BD with psychosis may be a biologically relevant subtype of BD. As these studies were focused on deep WM, future studies of SWM should be well-powered enabling categorization into homogenous subsamples.

Increased interstitial white matter neuron (IWMN) density has been reported in the SWM of the superior temporal gyrus ^9^ and prefrontal cortex in people with SZ ^6–8^. IWMNs, adult remnants of the embryonic cortical subplate ^63, 64^, reflect early developmental abnormalities and may contribute to our findings of abnormal gFA in the prefrontal cortex and superior temporal gyrus, and possibly other regions, in patients. Interneurons, facilitating communication between sensory and motor neurons, are fundamental for higher cognitive functions and are decreased in people with SZ ^65–67^. The inverse relationship found between increased IWMN in SWM and deficits in grey matter interneurons ^6^ suggest that SWM abnormalities are implicated in functional disconnectivity and subsequently cognitive impairment ^68, 69^. Overall, it has been suggested that decreased FA in SZ patients may be attributed to excessive synaptic pruning ^70^, which is supported by decreased spine density ^71, 72^ and presynaptic protein markers ^73^ found in the brains of people with SZ from postmortem studies. Increased FA in patients may reveal remodelling and growth of myelin in response to dysfunction, ^74^ thus serving as a structural compensation that may not be accompanied by benefits in function and behaviour.

### Limitations

This study is not without limitations. While some short association bundles have been validated by postmortem dissections ^75^, the current study defined bundles using tractography techniques ^21^, which may be prone to false positives. Nevertheless, we maximised reliability by only including bundles in our analyses that were constructed in all subjects, regardless of diagnosis.

Most patients were receiving antipsychotic medication and there are reports that first and second-generation antipsychotics are associated with both decreases ^76^ and increases ^77, 78^ in FA of WM. We found an inverse relationship between daily OLZ equivalent dose and gFA in six bundles after correction for multiple comparisons, suggesting that chronic antipsychotic use is associated with decreased SWM FA and may be a confounding variable. However, FA differences have also been reported in medication-naïve patients ^79^ and non-affected relatives of patients ^23^, suggesting that these disturbances represent a marker of the disease and are not solely due to medication effects. Our sample was not large enough to separate out individuals with BD taking lithium (n=13). There is evidence that lithium may increase FA in deep WM tracts associated with emotion processing in BD ^80, 81^, suggesting a normalization of WM microstructure following treatment. In this case, we would be less likely to detect differences between controls and BD patients; however, our findings were robust enough even if potentially influenced by effects of lithium given the significant findings. In line with the largest multi-site study to date examining deep WM in people with SZ ^82^, we found no relationship between duration of illness and gFA in either patient group when age was controlled for. Though, like most neuroimaging studies, our study was cross-sectional and therefore does not have the ability to determine whether our observed findings precede the onset of the disease or if they developed during its course.

The present study identified shared and distinct SWM abnormalities between SZ and BD. We demonstrate that these two disorders are not entirely distinct clinical entities at the level of SWM pathology, suggesting that their overlapping characteristics may be partially explained by common deviations found in SWM. These novel findings suggest that we should improve our understanding of the anatomy and function of short associative fibers in major psychiatric disorders which may be helpful in eventually preventing and mitigating the debilitating symptoms.

## Acknowledgements

We thank all subjects for their participation in this study. We would also like to thank Philippe Le Corvoisier for assisting in the recruitment of healthy controls.

## References

1. Bleuler E. Dementia praecox or the group of schizophrenias. Oxford, England: International Universities Press; 1950.

2. Friston KJ, Frith CD. Schizophrenia: a disconnection syndrome? Clinical neuroscience (New York, NY) 1995;3(2):89–97.

3. Davis KL, Stewart DG, Friedman JI, et al. White matter changes in schizophrenia: Evidence for myelin-related dysfunction. Archives of general psychiatry 2003;60(5):443–456.

4. Nortje G, Stein DJ, Radua J, Mataix-Cols D, Horn N. Systematic review and voxel-based meta-analysis of diffusion tensor imaging studies in bipolar disorder. J Affect Disord Sep 5 2013;150(2):192–200.

5. Phillips OR, Joshi SH, Squitieri F, et al. Major Superficial White Matter Abnormalities in Huntington’s Disease. Frontiers in Neuroscience 2016-May-23 2016;10(197).

6. Yang Y, Fung SJ, Rothwell A, Tianmei S, Weickert CS. Increased Interstitial White Matter Neuron Density in the Dorsolateral Prefrontal Cortex of People with Schizophrenia. Biological Psychiatry 1/1/ 2011;69(1):63–70.

7. Eastwood SL, Harrison PJ. Interstitial white matter neuron density in the dorsolateral prefrontal cortex and parahippocampal gyrus in schizophrenia. Schizophrenia research 11/15/2005;79(2-3):181–188.

8. Anderson SA, Volk DW, Lewis DA. Increased density of microtubule associated protein 2-immunoreactive neurons in the prefrontal white matter of schizophrenic subjects. Schizophrenia research 5// 1996;19(2-3):111–119.

9. Eastwood SL, Harrison PJ. Interstitial white matter neurons express less reelin and are abnormally distributed in schizophrenia: towards an integration of molecular and morphologic aspects of the neurodevelopmental hypothesis. Molecular psychiatry Sep 2003;8(9):769, 821–731.

10. Jones DK. The effect of gradient sampling schemes on measures derived from diffusion tensor MRI: a Monte Carlo study. Magnetic resonance in medicine Apr 2004;51(4):807–815.

11. Poonawalla AH, Zhou XJ. Analytical error propagation in diffusion anisotropy calculations. Journal of magnetic resonance imaging: JMRI Apr 2004;19(4):489–498.

12. Dong Q, Welsh RC, Chenevert TL, Carlos RC, Maly-Sundgren P, Gomez-Hassan DM, Mukherji SK. Clinical applications of diffusion tensor imaging. Journal of magnetic resonance imaging: JMRI Jan 2004;19(1):6–18.

13. Neil JJ, Shiran SI, McKinstry RC, et al. Normal brain in human newborns: apparent diffusion coefficient and diffusion anisotropy measured by using diffusion tensor MR imaging. Radiology Oct 1998;209(1):57–66.

14. Song SK, Sun SW, Ju WK, Lin SJ, Cross AH, Neufeld AH. Diffusion tensor imaging detects and differentiates axon and myelin degeneration in mouse optic nerve after retinal ischemia. NeuroImage Nov 2003;20(3):1714–1722.

15. Takahashi M, Hackney DB, Zhang G, et al. Magnetic resonance microimaging of intraaxonal water diffusion in live excised lamprey spinal cord. Proceedings of the National Academy of Sciences of the United States of America Dec 10 2002;99(25):16192–16196.

16. Takahashi M, Ono J, Harada K, Maeda M, Hackney DB. Diffusional anisotropy in cranial nerves with maturation: quantitative evaluation with diffusion MR imaging in rats. Radiology Sep 2000;216(3):881–885.

17. Buchsbaum MS, Schoenknecht P, Torosjan Y, et al. Diffusion tensor imaging of frontal lobe white matter tracts in schizophrenia. Annals of General Psychiatry 11/28 02/17/received 11/28/accepted 2006;5:19–19.

18. Mori T, Ohnishi T, Hashimoto R, et al. Progressive changes of white matter integrity in schizophrenia revealed by diffusion tensor imaging. Psychiatry research Feb 28 2007;154(2):133–145.

19. Ashtari M, Cottone J, Ardekani BA, Cervellione K, Szeszko PR, Wu J, Chen S, Kumra S. Disruption of white matter integrity in the inferior longitudinal fasciculus in adolescents with schizophrenia as revealed by fiber tractography. Archives of general psychiatry Nov 2007;64(11):1270–1280.

20. Szeszko PR, Robinson DG, Ashtari M, et al. Clinical and neuropsychological correlates of white matter abnormalities in recent onset schizophrenia. Neuropsychopharmacology: official publication of the American College of Neuropsychopharmacology Apr 2008;33(5):976–984.

21. Guevara M, Roman C, Houenou J, Duclap D, Poupon C, Mangin JF, Guevara P. Reproducibility of superficial white matter tracts using diffusion-weighted imaging tractography. NeuroImage Feb 15 2017;147:703–725.

22. Nazeri A, Chakravarty MM, Felsky D, Lobaugh NJ, Rajji TK, Mulsant BH, Voineskos AN. Alterations of superficial white matter in schizophrenia and relationship to cognitive performance. Neuropsychopharmacology: official publication of the American College of Neuropsychopharmacology Sep 2013;38(10):1954–1962.

23. Phillips OR, Nuechterlein KH, Asarnow RF, et al. Mapping corticocortical structural integrity in schizophrenia and effects of genetic liability. Biol Psychiatry Oct 1 2011;70(7):680–689.

24. Cabeen RP, Laidlaw DH, Ruggieri A, Dickstein DP. Preliminary mapping of the structural effects of age in pediatric bipolar disorder with multimodal MR imaging. Psychiatry Research: Neuroimaging 2018/03/30/ 2018;273:54–62.

25. Zhang S, Wang Y, Deng F, et al. Disruption of superficial white matter in the emotion regulation network in bipolar disorder. NeuroImage Clinical Sep 26 2018;20:875–882.

26. Gandal MJ, Haney JR, Parikshak NN, et al. Shared molecular neuropathology across major psychiatric disorders parallels polygenic overlap. Science 2018;359(6376):693–697.

27. Nazeri A, Chakravarty MM, Rajji TK, et al. Superficial white matter as a novel substrate of age-related cognitive decline. Neurobiology of aging 6// 2015;36(6):2094–2106.

28. Katz J, d’Albis MA, Boisgontier J, et al. Similar white matter but opposite grey matter changes in schizophrenia and high-functioning autism. Acta Psychiatrica Scandinavica 2016;In Press.

29. Sarrazin S, Poupon C, Linke J, et al. A multicenter tractography study of deep white matter tracts in bipolar I disorder: psychotic features and interhemispheric disconnectivity. JAMA psychiatry Apr 2014;71(4):388–396.

30. Tuch DS, Reese TG, Wiegell MR, Wedeen VJ. Diffusion MRI of complex neural architecture. Neuron Dec 4 2003;40(5):885–895.

31. Woods SW. Chlorpromazine equivalent doses for the newer atypical antipsychotics. The Journal of clinical psychiatry Jun 2003;64(6):663–667.

32. Benjamini Y, Hochberg Y. Controlling the False Discovery Rate: A Practical and Powerful Approach to Multiple Testing. Journal of the Royal Statistical Society Series B (Methodological) 1995;57(1):289–300.

33. Cohen J. Statistical Power Analysis for the Behavioral Sciences: Academic Press; 1969.

34. Ioannidis JPA, Munafò MR, Fusar-Poli P, Nosek BA, David SP. Publication and other reporting biases in cognitive sciences: detection, prevalence and prevention. Trends in cognitive sciences 03/18 2014;18(5):235–241.

35. Chrobak AA, Siuda-Krzywicka K, Siwek GP, Tereszko A, Janeczko W, Starowicz-Filip A, Siwek M, Dudek D. Disrupted implicit motor sequence learning in schizophrenia and bipolar disorder revealed with ambidextrous Serial Reaction Time Task. Progress in neuro-psychopharmacology & biological psychiatry Oct 3 2017;79(Pt B):169–175.

36. Hill SK, Harris MSH, Herbener ES, Pavuluri M, Sweeney JA. Neurocognitive Allied Phenotypes for Schizophrenia and Bipolar Disorder. Schizophrenia Bulletin 2008;34(4):743–759.

37. Cohen AS, McGovern JE, Dinzeo TJ, Covington MA. Speech Deficits in Serious mental Illness: A Cognitive Resource Issue? Schizophrenia research 11/17 2014;160(0):173–179.

38. Oh A, Duerden EG, Pang EW. The role of the insula in speech and language processing. Brain and language Aug 2014;135:96–103.

39. Morice R, McNicol D. Language changes in schizophrenia: a limited replication. Schizophr Bull 1986;12(2):239–251.

40. Raucher-Chene D, Achim AM, Kaladjian A, Besche-Richard C. Verbal fluency in bipolar disorders: A systematic review and meta-analysis. J Affect Disord Jan 1 2017;207:359–366.

41. Harvey PD. Mood Symptoms, Cognition, and Everyday Functioning: in Major Depression, Bipolar Disorder, and Schizophrenia. Innovations in Clinical Neuroscience 10/ 2011;8(10):14–18.

42. Carter CS, Botvinick MM, Cohen JD. The contribution of the anterior cingulate cortex to executive processes in cognition. Reviews in the neurosciences 1999;10(1):49–57.

43. Vöhringer P, Barroilhet S, Amerio A, Reale M, Vergne D, Alvear K, Ghaemi S. Cognitive Impairment in Bipolar Disorder and Schizophrenia: A Systematic Review. Frontiers in psychiatry 2013-August-08 2013;4(87).

44. Barta PE, Pearlson GD, Powers RE, Richards SS, Tune LE. Auditory hallucinations and smaller superior temporal gyral volume in schizophrenia. The American journal of psychiatry Nov 1990;147(11):1457–1462.

45. Rajarethinam RP, DeQuardo JR, Nalepa R, Tandon R. Superior temporal gyrus in schizophrenia: a volumetric magnetic resonance imaging study. Schizophrenia research Jan 21 2000;41(2):303–312.

46. John JP. Fronto-temporal dysfunction in schizophrenia: A selective review. Indian Journal of Psychiatry Jul-Sep 2009;51(3):180–190.

47. Lohr JB, Caligiuri MP. Lateralized hemispheric dysfunction in the major psychotic disorders: historical perspectives and findings from a study of motor asymmetry in older patients. Schizophrenia research Oct 30 1997;27(2-3):191–198.

48. Love S. Demyelinating diseases. Journal of Clinical Pathology 09/26/accepted 2006;59(11):1151–1159.

49. Mulert C, Kirsch V, Whitford TJ, et al. Hearing voices: a role of interhemispheric auditory connectivity? The world journal of biological psychiatry: the official journal of the World Federation of Societies of Biological Psychiatry Feb 2012;13(2):153–158.

50. Hubl D, Koenig T, Strik W, et al. Pathways that make voices: white matter changes in auditory hallucinations. Archives of general psychiatry Jul 2004;61(7):658–668.

51. Seok JH, Park HJ, Chun JW, Lee SK, Cho HS, Kwon JS, Kim JJ. White matter abnormalities associated with auditory hallucinations in schizophrenia: a combined study of voxel-based analyses of diffusion tensor imaging and structural magnetic resonance imaging. Psychiatry research Nov 15 2007;156(2):93–104.

52. Shergill SS, Kanaan RA, Chitnis XA, et al. A diffusion tensor imaging study of fasciculi in schizophrenia. The American journal of psychiatry Mar 2007;164(3):467–473.

53. Garrity AG, Pearlson GD, McKiernan K, Lloyd D, Kiehl KA, Calhoun VD. Aberrant “default mode” functional connectivity in schizophrenia. The American journal of psychiatry Mar 2007;164(3):450–457.

54. Zhou Y, Liang M, Tian L, Wang K, Hao Y, Liu H, Liu Z, Jiang T. Functional disintegration in paranoid schizophrenia using resting-state fMRI. Schizophrenia research Dec 2007;97(1-3):194–205.

55. Guo W, Liu F, Chen J, Wu R, Li L, Zhang Z, Chen H, Zhao J. Hyperactivity of the default-mode network in first-episode, drug-naive schizophrenia at rest revealed by family-based case–control and traditional case–control designs. Medicine 03/31

56. Zhou L, Pu W, Wang J, et al. Inefficient DMN Suppression in Schizophrenia Patients with Impaired Cognitive Function but not Patients with Preserved Cognitive Function. Scientific Reports 02/17/online 2016;6:21657.

57. Northoff G, Heinzel A, de Greck M, Bermpohl F, Dobrowolny H, Panksepp J. Self-referential processing in our brain--a meta-analysis of imaging studies on the self. NeuroImage May 15 2006;31(1):440–457.

58. Smith SM, Jenkinson M, Johansen-Berg H, et al. Tract-based spatial statistics: Voxelwise analysis of multi-subject diffusion data. NeuroImage 2006/07/15/ 2006;31(4):1487–1505.

59. Ouyang M, Kang H, Detre JA, Roberts TPL, Huang H. Short-range connections in the developmental connectome during typical and atypical brain maturation. Neuroscience and biobehavioral reviews Dec 2017;83:109–122.

60. Barysheva M, Jahanshad N, Foland-Ross L, Altshuler LL, Thompson PM. White matter microstructural abnormalities in bipolar disorder: A whole brain diffusion tensor imaging study. NeuroImage: Clinical 2013/01/01/ 2013;2:558–568.

61. Sussmann JE, Lymer GK, McKirdy J, et al. White matter abnormalities in bipolar disorder and schizophrenia detected using diffusion tensor magnetic resonance imaging. Bipolar disorders Feb 2009;11(1):11–18.

62. Skudlarski P, Schretlen DJ, Thaker GK, et al. Diffusion Tensor Imaging White Matter Endophenotypes in Patients With Schizophrenia or Psychotic Bipolar Disorder and Their Relatives. American Journal of Psychiatry 2013/08/01 2013;170(8):886–898.

63. Kostovic I, Rakic P. Cytology and time of origin of interstitial neurons in the white matter in infant and adult human and monkey telencephalon. Journal of neurocytology Apr 1980;9(2):219–242.

64. Chun JJ, Shatz CJ. Interstitial cells of the adult neocortical white matter are the remnant of the early generated subplate neuron population. The Journal of comparative neurology Apr 22 1989;282(4):555–569.

65. Hashimoto T, Volk DW, Eggan SM, Mirnics K, Pierri JN, Sun Z, Sampson AR, Lewis DA. Gene expression deficits in a subclass of GABA neurons in the prefrontal cortex of subjects with schizophrenia. The Journal of neuroscience: the official journal of the Society for Neuroscience Jul 16 2003;23(15):6315–6326.

66. Guidotti A, Auta J, Davis JM, et al. Decrease in reelin and glutamic acid decarboxylase67 (GAD67) expression in schizophrenia and bipolar disorder: a postmortem brain study. Archives of general psychiatry Nov 2000;57(11):1061–1069.

67. Akbarian S, Kim JJ, Potkin SG, Hagman JO, Tafazzoli A, Bunney WE, Jr., Jones EG. Gene expression for glutamic acid decarboxylase is reduced without loss of neurons in prefrontal cortex of schizophrenics. Archives of general psychiatry Apr 1995;52(4):258–266.

68. Lodge DJ, Behrens MM, Grace AA. A loss of parvalbumin-containing interneurons is associated with diminished oscillatory activity in an animal model of schizophrenia. The Journal of neuroscience: the official journal of the Society for Neuroscience 2009;29(8):2344–2354.

69. Murray AJ, Woloszynowska-Fraser MU, Ansel-Bollepalli L, Cole KLH, Foggetti A, Crouch B, Riedel G, Wulff P. Parvalbumin-positive interneurons of the prefrontal cortex support working memory and cognitive flexibility. Scientific Reports 11/26/online 2015;5:16778.

70. Feinberg I. Schizophrenia: Caused by a fault in programmed synaptic elimination during adolescence? Journal of psychiatric research 1982/01/01/ 1982;17(4):319–334.

71. Glantz LA, Lewis DA. Decreased dendritic spine density on prefrontal cortical pyramidal neurons in schizophrenia. Archives of general psychiatry Jan 2000;57(1):65–73.

72. Broadbelt K, Byne W, Jones LB. Evidence for a decrease in basilar dendrites of pyramidal cells in schizophrenic medial prefrontal cortex. Schizophrenia research Nov 1 2002;58(1):75–81.

73. Faludi G, Mirnics K. Synaptic changes in the brain of subjects with schizophrenia. International journal of developmental neuroscience: the official journal of the International Society for Developmental Neuroscience May 2011;29(3):305–309.

74. Palaniyappan L. Progressive cortical reorganisation: A framework for investigating structural changes in schizophrenia. Neuroscience and biobehavioral reviews Aug 2017;79:1–13.

75. Vergani F, Mahmood S, Morris CM, Mitchell P, Forkel SJ. Intralobar fibres of the occipital lobe: A post mortem dissection study. Cortex 2014/07/01/ 2014;56:145–156.

76. Wang Q, Cheung C, Deng W, et al. White-matter microstructure in previously drug-naive patients with schizophrenia after 6 weeks of treatment. Psychological medicine Nov 2013;43(11):2301–2309.

77. Xiao L, Xu H, Zhang Y, et al. Quetiapine facilitates oligodendrocyte development and prevents mice from myelin breakdown and behavioral changes. Molecular psychiatry Jul 2008;13(7):697–708.

78. Ozcelik-Eroglu E, Ertugrul A, Oguz KK, Has AC, Karahan S, Yazici MK. Effect of clozapine on white matter integrity in patients with schizophrenia: a diffusion tensor imaging study. Psychiatry research Sep 30 2014;223(3):226–235.

79. Cheung V, Cheung C, McAlonan GM, et al. A diffusion tensor imaging study of structural dysconnectivity in never-medicated, first-episode schizophrenia. Psychological medicine 2007;38(6):877–885.

80. Kafantaris V, Spritzer L, Doshi V, Saito E, Szeszko PR. Changes in white matter microstructure predict lithium response in adolescents with bipolar disorder. Bipolar disorders Nov 2017;19(7):587–594.

81. Benedetti F, Absinta M, Rocca MA, et al. Tract-specific white matter structural disruption in patients with bipolar disorder. Bipolar disorders Jun 2011;13(4):414–424.

82. Kay SR, Fiszbein A, Opler LA. The positive and negative syndrome scale (PANSS) for schizophrenia. Schizophr Bull 1987;13(2):261–276.

